# Resuscitation of soil microbiota after > 70-years of desiccation

**DOI:** 10.1101/2020.11.06.371641

**Authors:** Jun Zhao, Dongfeng Chen, Wei Gao, Zhiying Guo, Zhongjun Jia, Marcela Hernández

## Abstract

The abundance and diversity of bacteria in 24 historical soil samples under air-dried storage conditions for more than 70 years were assessed by quantification and high-throughput sequencing analysis of 16S rRNA genes. All soils contained a measurable abundance of bacteria varying from 10^3^ to 10^8^ per gram of soil and contrasting community compositions were observed in different background soils, suggesting that the bacteria detected were indigenous to the soil. Following a 4-week soil rewetting event, the bacterial abundance significantly increased in soils, indicating strong adaptation of soil bacteria to extreme osmotic change and high resuscitation potential of some bacteria over long periods of desiccation. *Paenibacillus, Cohnella* and two unclassified *Bacillales* genera within the phylum *Firmicutes* represented the most ubiquitously active taxa, which showed growth in the highest number of soils (≥12 soils), while genera *Tumebacillus, Alicyclobacillus* and *Brevibacillus* in the phylum *Firmicutes* displayed the highest growth rates in soils (with >1000-fold average increase) following rewetting. Additionally, some *Actinobacteria* and *Proteobacteria* genera showed relatively high activity following rewetting, suggesting that the resilience to long-term desiccation and rewetting is a common trait across phylogenetically divergent microbes. The present study thus demonstrated that diversified groups of microbes are present and potentially active in historically desiccated soils, which might be of importance in the context of microbial ecology.

## 1 Introduction

Although frozen storage is widely accepted as the optimal method for preserving the chemical and biological components of soils, desiccation (i.e. air-drying) is more commonly used for large scale or cross-generational soil archives [1]. The desiccated soils, even after decades of storage, have been frequently used for analyzing the chemical composition [2–6], but the persistence of the biological components, especially the microbes in these soils remain largely untested. Microbes are famously known for tolerance to extreme environmental stresses, exemplified by growth of a few isolated strains after millions of years of dormancy under extremely low thermal conditions [7–10]. In respect to water stress, microbes have demonstrated strong resistance to long periods of drought based on both the cultivation of pure strains and the ecological investigation of samples from arid ecosystems [11, 12]. For instance, taxonomically diverse groups of bacteria are abundantly present and detectable in perennial drylands, including members from phyla of *Actinobacteria*, *Bacteroidetes* and *Proteobacteria* [13–15], suggesting that the ability of adaptation to long periods of drought is not restricted to a few particular taxa [16]. Compared to natural arid soils, the desiccated soils systematically archived in the laboratory have endured persistent water depletion without intermittent rain precipitation as occurs in the field. The careful curation of the soils in the laboratory prevents the occurrence of other major environmental perturbations, such as vegetation and animal activities. Additionally, the air-dried soil samples can also provide information on drought tolerance of microbial communities that originated from non-arid ecosystems. Previous studies on historical samples after decades of storage in air-dried conditions has indeed identified different groups of bacterial and eukaryotic phylotypes, and suggested the possibility to use these samples to detect systematic differences in soil microbial community composition [17–22]. However, due to the low depth of sequencing, the information on microbial abundance and diversity revealed from these studies is limited.

Water availability is a key factor controlling soil microbial activity and regulating terrestrial ecosystem functioning. A decrease in soil moisture following desiccation can cause high osmotic stress to the cells and prevent the diffusion of soluble nutrient substrates [23], leading to dormancy of many soil microbes and subsequent decline in many important biogeochemical processes including nitrification [24], denitrification [25, 26] and methanotrophy [27, 28]. Some soil microbes can be reactivated soon after the alleviation of a short period of drought stress, usually following a rewetting event [24, 29–31]. Different phylotypes of microbes appear to respond differentially to rewetting, with their abundance recovered at different rates from within one hour to after a few days [32]. However, it is not known how different taxa respond to a rewet event following a long-term, continuous desiccation.

In the present study, we acquired 24 historical soil samples collected from across mainland China during the years of 1934-1939 from the Soil Archives at the Institute of Soil Science, Chinese Academy of Sciences. By assessing the abundances (by quantitative PCR of 16S rRNA genes) and community compositions (by pyrosequencing) of bacteria in desiccated condition and after a rewet incubation, our study aims to assess the resuscitation potentials of microbes through decades of desiccation and identify the most responsive taxa in these soils.

## 2 Material and methods

### 2.1 Soil description and preparation

Samples were collected from seven provinces across the mainland of China during the years of 1934-1939 (Fig. S1), and stored in air-dried conditions. The available records by the collectors indicate widely spread field origins and/or properties for most of the soils, including different locations, soil taxonomies and vegetation (Table 1). Soon after each collection, a small part of the soil was kept in a reservation box in air-dried conditions as part of the soil archives at the Institute of Soil Science, Chinese Academy of Sciences. The weight of the archive soil was about 15-50 g for each sample and completely dried by visual inspection. Due to the scarcity in quantity and historic values of the samples, we obtained no more than 10 g of each soil for following molecular analysis and did not perform tests on the soil chemical and physical properties.

**Table 1.** The documented information of archive soils.

| Sample No. | Series No. | Collecting date | Collector(s) | Province | City/County | Land use | Vegetation | Soil genetic classification | Soil parent materials | Soil taxonomy |
| --- | --- | --- | --- | --- | --- | --- | --- | --- | --- | --- |
| S01 | 5151 | 1934/10/9 | Po Suo, Guangjiong Hou | Qinghai | Dulan | livestock farming | - | calcareous clay | - | uap-ustic isohumisols |
| S02 | 5213 | 1935/12/24 | Lianqing Zhu et. al | Sichuan | Huayang | dry land | - | brown soil | granite | hapli-udic argosols |
| S03 | 5009 | 1939/11/15 | Lianqing Zhu, Fanqi Zeng | Yunnan | Mengzi | farmland | rice | red soil | - | Fe-leachi-stagnic anthrosols |
| S04 | 5157 | 1934/11/13 | Lianjie Li et. al | Guangxi | Yongning | wasteland | miscellaneous tree e.g. Chinese fir, pine | red soil | sandstone | hap-udic ferrisols |
| S05 | HN | 1935/1/1 | Changyun Zhou | Hunan | Changsha | - | - | red soil | - | - |
| S06 | 5001 | 1934/11/13 | Guangjiong Hou | Jiangxi | Nanchang | - | - | red soil | - | hap-udic ferrisols |
| S07 | 5289 | 1938/5/24 | Daquan Song et. al | Fujian | Jianyang | vegetable garden | vegetables | fluvisols | river alluvium | acidic humid alluvial entisols |
| S08 | 5143 | 1938/10/1 | Daquan Song et. al | Fujian | Longyan | wasteland | - | lime soil | limestone | black lithologic isohumisols |
| S09 | 5277 | 1938/5/24 | Daquan Song et. al | Fujian | Chongan | tea orchard | tea | purple soil | calcareous purple shale | calcareous purplish-udic cambosols |
| S10 | 5061 | 1938/12/1 | Daquan Song et. al | Fujian | Jianyang | wasteland | - | podzol | - | typic spodosol |
| S11 | 5062 | 1938/12/1 | Daquan Song et. al | Fujian | Dehua | wasteland | grass | podzol | volcanic rock | typic spodosol |
| S12 | 5275 | 1938/5/3 | Daquan Song et. al | Fujian | Chongan | wasteland | weeds | purple soil | purple sandshale | purplish-udic cambosols |
| S13 | 5030 | 1938/10/23 | Daquan Song, Zhenyu Yu | Fujian | Longxi | farmland | rice | red soil | river alluvium | typic Fe-accumulic-stagnic anthrosols |
| S14 | 5031 | 1938/10/23 | Daquan Song et. al | Fujian | Jianou | farmland | rice | red soil | river alluvium | typic Fe-accumulic-stagnic anthrosols |
| S15 | 5032 | 1938/12/23 | Daquan Song et. al | Fujian | Longxi | farmland | rice | red soil | - | typic Fe-accumulic-stagnic anthrosols |
| S16 | 5033 | 1938/12/23 | Daquan Song et. al | Fujian | Longxi | farmland | rice | red soil | - | Fe-accumulic-stagnic anthrosols |
| S17 | 5171 | 1938/8/5 | Daquan Song et. al | Fujian | Yongchun | wasteland | shrubs and weeds | red soil | white effusive rock | hap-udic ferrisols |
| S18 | 5172 | 1938/5/5 | Daquan Song et. al | Fujian | Yongchun | wasteland | evergreen broad-leaved trees | red soil | quartzite and sandstone | humic udic ferrisols |
| S19 | 5174 | 1938/5/5 | Daquan Song et. al | Fujian | Yongchun | wasteland | evergreen broad-leaved | red soil | quartzite | hap-udic ferrisols |
| S20 | 5274 | 1938/10/3 | Changyun Zhou et. al | Fujian | Yongchun | wasteland | grass | acidic residual soil | granite | acidic ochreous-aquic cambosols |
| S21 | 5278 | 1938/5/24 | Changyun Zhou et. al | Fujian | Yongan | wasteland | - | red soil | sandy lake sediment | semi-humic organic soil |
| S22 | 5119 | 1938/10/22 | Chengfan Xi et. al | Fujian | Putian | wasteland | natural vegetation | coastal saline soil | sea mud alluvium | halogenic soil |
| S23 | 5120 | 1938/10/22 | Chengfan Xi et. al | Fujian | Putian | wasteland | natural vegetation | coastal saline soil | - | halogenic soil |
| S24 | 5173 | 1938/5/5 | Daquan Song et. al | Fujian | Zhangping | wasteland | weeds | yellow soil | granite | humic udic ferrisols |

### 2.2 Rewetting microcosm incubation

For each microcosm, 1 g of soils were added into sterilized 5-ml centrifuge tubes and rewetted up to 30% of water holding capacity with sterile water. The tubes were sealed and incubated at 28°C for 28 days. Soil samples were destructively collected and frozen at −80°C for molecular analyses. All soils were subjected to rewetting microcosm incubations in triplicate, except for S01, S05, S23 and S24, which were incubated in duplicate due to a low soil quantity obtained.

### 2.3 DNA extraction

DNA was extracted from 1 g of soil, using FastDNA spin kit for DNA extraction (MP Biomedicals, USA), according to the manufacturer’s instructions, and dissolved in a final volume of 50 μl sterile water. The quantity and quality of DNA extracts were measured using a NanoDrop ND-1000 UV-visible light spectrophotometer (NanoDrop Technologies, USA).

### 2.4 Real-time quantitative PCR

Real-time quantitative PCR (qPCR) was conducted on a CFX96 Optical Real-Time detection system (Bio-Rad Laboratories, Inc., Hercules, CA, USA) to determine the abundance of 16S rRNA genes in total DNA extracts, using universal PCR primers 515F (GTGCCAGCMGCCGCGG) and 907R (CCGTCAATTCMTTTRAGTTT) [33]. The thermal process started with 3 min of incubation at 95°C, followed by 40 cycles of 95°C for 30s, 55°C for 30s, and 72°C for 30s with plate read, and ended with a melt curve from 65 to 95°C with increments of 0.5°C per 5s. The real-time quantitative PCR standards were generated using plasmid DNA from one representative clone containing 16S rRNA genes, and a dilution series of standard template over eight orders of magnitude per assay was used. Amplification efficiencies ranged from 96% to 105%, with R^2^ values of 0.992 to 0.996.

### 2.5 High-throughput sequencing and analysis

High-throughput sequencing was performed to investigate the bacterial community composition in soils. 16S rRNA gene fragments were amplified using the same universal 515F-907R primer pair but with an adaptor, key sequence, and unique tag sequence for each sample. The thermal conditions were as follows: 5 min of incubation at 94°C, followed by 30 cycles of 94°C for 30s, 54°C for 30s, and 72°C for 45s, with a 10 min extension at 72°C. The resulting PCR products were gel purified and combined in equimolar ratios into single tubes before pyrosequencing. The pyrosequencing was conducted on a Roche 454 GS FLX Titanium sequencer (Roche Diagnostics Corporation, Bran-ford, CT, USA).

All sequencing data were processed using multiple software packages. Reads from each sample were first sorted by specific tag sequences using Mothur software [34]. Sequences were further trimmed, and only reads of > 350 bp in length with an average quality score > 25 and no ambiguous base calls were included for subsequent analyses. The operational taxonomic units (OTUs) were clustered at 97% similarity cutoff by UPARSE algorithm after the removal of chimeras using QIIME software [35]. A representative sequence of each OTU was classified using the SILVA-132 16S rRNA gene database (bootstrap confidence threshold of 80 %) in Mothur [34].

The raw sequencing data of all samples have been deposited in the European Nucleotide Archive (ENA) with accession numbers ERR1520684 and ERR1520685.

### 2.6 Statistical analyses

The absolute abundance of total bacteria or a specific taxon in each soil microcosm was calculated by multiplying the total 16S rRNA gene number (results from qPCR) by the relative abundance of bacteria or the taxon (results of amplicon sequencing). The nonparametric Wilcoxon test was used to determine the significance in difference in bacterial abundance between all the dry and rewet soils. The fold change in abundance of total bacteria or a specific taxon in each soil was further calculated as the ratio of abundance after rewetting to that in the desiccated condition. The abundance of a taxon was considered to increase in the soil when the fold change was significantly higher than 1 following a t-test (*p* < 0.05), and the corresponding taxon was defined as active.

Heatmaps were constructed as previously described in Hernández et al., [36] using the vegan package in R. The samples and OTUs were clustered according to Euclidean distances between all Hellinger transformed data.

## 3 Results

### 3.1 Change in bacterial abundance

Total bacterial abundance was calculated based on the qPCR and high-throughput sequencing of 16S rRNA genes. The 28-day rewet incubation led to significant increase in bacterial abundance in most soils (n=24) (Fig. 1a). The total bacterial abundances in soils were 1.9×10^3^ to 1.7×10^8^ per gram of dry weight soil (g^−1^ *d.w.s*), and reached 2.6×10^3^ to 4.1×10^8^ g^−1^ *d.w.s* following the rewetting incubation (Fig. S2). Specifically, bacterial abundance showed significant increase in 18 soils (*P* < 0.05), remained unchanged in 5 soils and only in 1 soil (S21) declined following the rewet. Notably, bacterial abundance in two soils (S03 and S05) increased by > 100-fold and in another seven soils (S04, S06, S08, S09, S12, S14 and S16) increased by > 10-fold following the rewetting incubation (Fig. 1b). These results demonstrated that some bacterial communities had high growth rates after regaining water in the soils. However, we cannot rule out the possibility that a large portion of the detected genes in the dry soils were residues from already dead cells, i.e. extracellular DNA [37, 38]. These cell residues could potentially become accessible nutrients for the living cells in the soils following rewetting [39], which might explain the decreased total bacterial abundance in one of the soils (S21) after the incubation.

**Figure 1.**
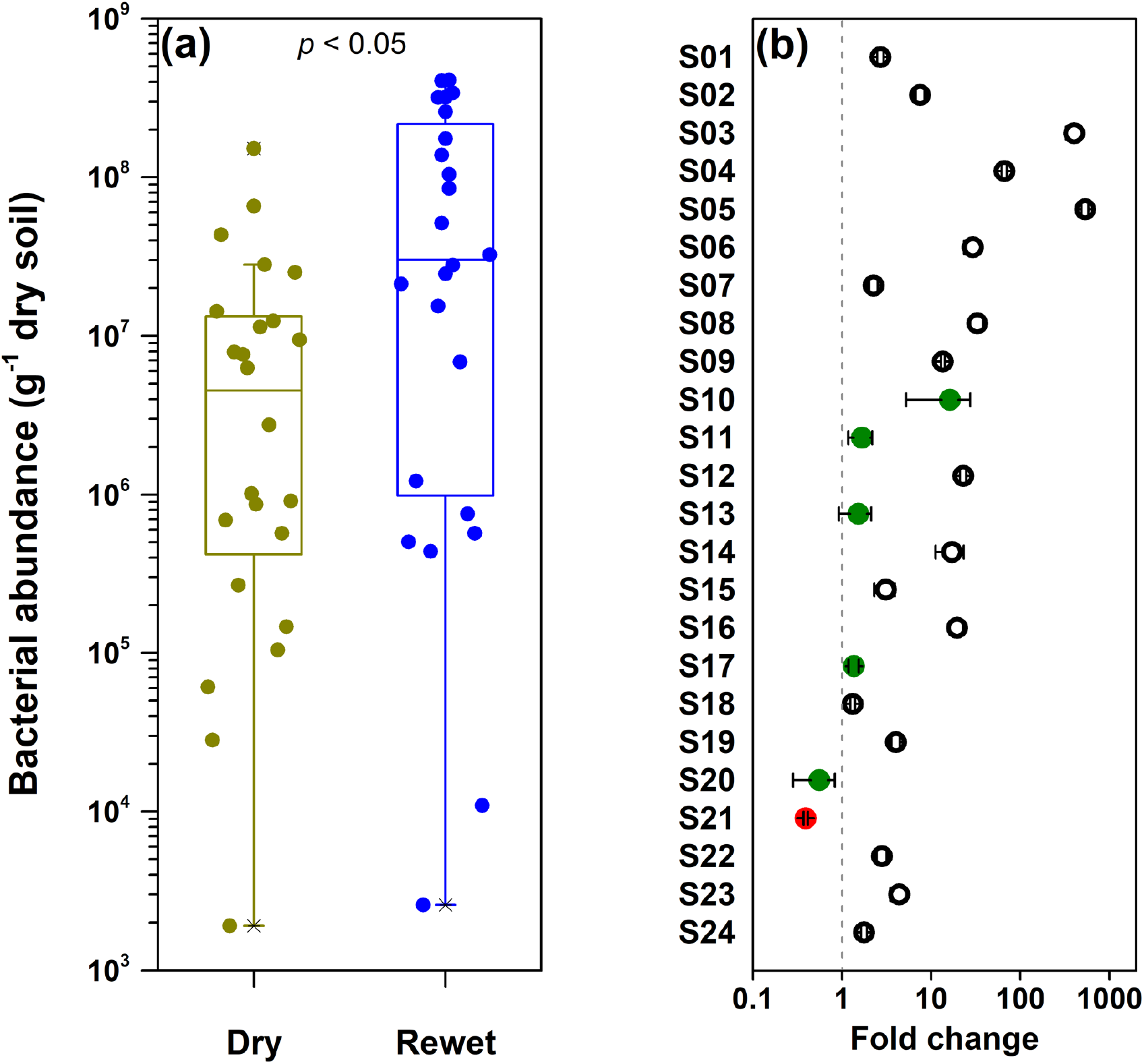
(a) Box-plot of bacterial abundance in soils in desiccated conditions (Dry) and after a rewetting incubation (Rewet). Centre lines show median value and boxes show upper and lower quartiles. Nonparametric Wilcoxon test was used to determine the significant difference of bacterial abundance in soils and *p* < 0.05 indicates significant difference between dry and rewetted soils (n=24 for both conditions). (b) The fold change of bacterial abundance following the rewetting incubation compared to the desiccated soils. The open black cycle indicates a fold change significantly greater than 1 (*p* < 0.05), the solid green circles represent no significant difference from 1 (*p* > 0.05), and the solid red circle indicates a change significantly lower than 1 (*P* < 0.05).

### 3.2 Change in bacterial composition

Taxonomic classification of 16S rRNA genes revealed a broad spectrum of bacterial taxa in soils. A total of 32 bacterial phyla were identified in soils at the phylum level. In the desiccated soils, the most abundant bacterial phyla identified were *Firmicutes* (averagely 55.4% of total bacteria), *Actinobacteria* (19.9%), *Proteobacteria* (9.7%), *Chloroflexi* (6.3%), *Acidobacteria* (1.7%) and *Planctomycetes* (1.1%) (Fig. 2 and S3a). Following the rewetting incubation, *Firmicutes* showed the highest growth rate and became the most abundant phylum in soils (80.4%), followed by *Proteobacteria* (9.4%), *Actinobacteria* (3.5%), *Acidobacteria* (1.6%) and *Chloroflexi* (1.0%) (Fig. 2 and S3b).

**Figure 2.**
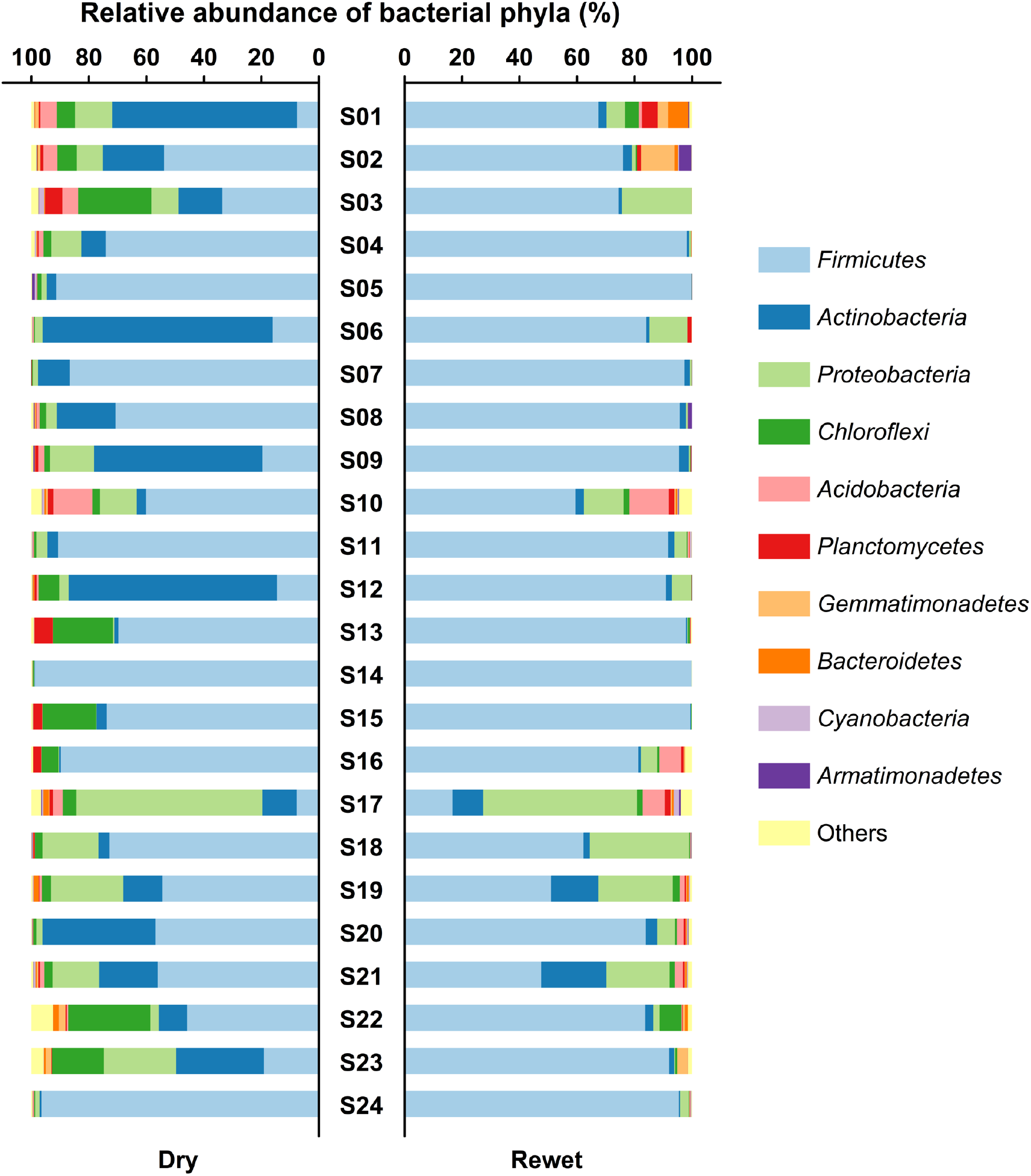
Composition of the bacteria in soils under desiccated conditions (Dry) and after the rewetting incubation (Rewet). The bacterial composition was plotted at phylum level and the most abundant phyla (with relative abundance >1% in average) are shown. Taxa not seen more than three times in at least 20 % of the samples were removed using phyloseq package in *R*. The relative abundances of minor phyla or classes were summed up and shown as “others” which do not exceed 7.5% of total bacterial reads in any given soil.

### 3.3 Identification active taxa

The activity of a specific bacterial taxon in each soil was assessed by the fold change in abundance after the rewetting incubation. At the genus level, a total of 759 taxa were identified in all soils, and 183 of them showed significant increase in abundance in at least one soil following the rewet (with fold change > 1 relative to the desiccated condition). Fig. 3 shows the top 21 genera with significant increases in abundance in more than three of the soils. *Paenibacillus*, *Cohnella* and two other unclassified *Bacillales* genera represented the most ubiquitously activated genera in soils, which showed increases in at least half of the soils following rewetting incubations (Fig. 3). On the other hand, *Bacillales* genera *Tumebacillus*, *Alicyclobacillus* and *Brevibacillus* showed the highest growth rate following rewetting, with on average >1,000-fold increase in abundance in these soils (Fig. 3). Although *Bacillales* comprised the most active microbial communities in our soils, other bacterial taxa, represented by one unclassified *Actinobacteria* genus, two *Gammaproteobacteria* genera (*Halomonas* and *Ralstonia*) and four genera belonging to *Firmicutes* order *Clostridia*, also showed marked growth in soils (Fig. 3).

**Figure 3.**
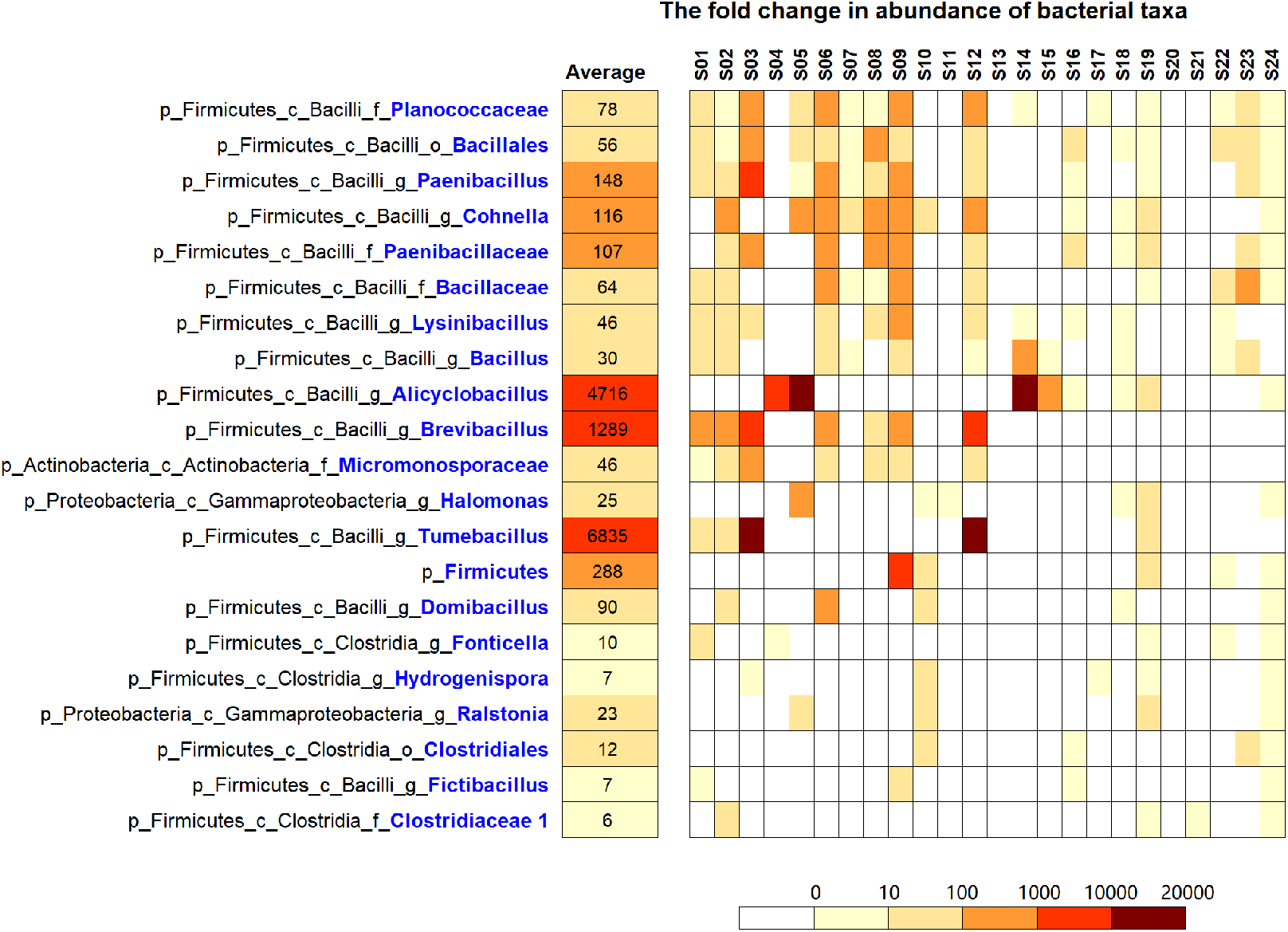
Heatmap showing significant fold change (*p* < 0.05) in abundance of active taxa in different soils following the 28-day rewetting incubation. A total of 21 taxa at the genus level are listed that displayed growth in the highest number of soils (> 3 soils). Each taxa designation contains the phylum (p), class (c) and genus (g) affiliation. Whenever a genus designation is not defined (unclassified genus), a defined family (f) or order (o) affiliation is specified instead. The values in the column “Average” indicates the average fold change in abundance of each taxa in soils following rewetting. Blank (white) square indicates no significant increase in abundance in a soil after rewetting (*p* > 0.05).

## Discussion

Our study described both the quantitative and compositional patterns of soil bacterial communities after continuous desiccation for decades. Phylogenetically diverse guilds of bacteria with measurable abundance were identified in these historical soils (Fig. 2 and S3a), demonstrating high resistance to the long-term desiccation. Driven by the stress from both water and nutrient deprivation for more than seven decades, the bacteria abundantly detected in the dry soils were most likely in a dormant state [23]. This was supported by the dominant presence of *Firmicutes* in most of the desiccated soils (Fig. 2), which consisted of *Paenibacillus*, *Cohnella, Bacillus* and *Clostridium* species (Fig. S3a) that are well known for producing resting bodies of endospores. In addition, in many soils, the frequent presence of taxa from *Actinobacteria*, *Proteobacteria* and *Chloroflexi* (Fig. S3a), which are less known for tolerance to extreme drought stress, implies the ubiquity of dormant life strategies against long-term desiccation across phylogenetically diverse microbial populations [16, 40]. This might be important in the maintenance of microbial biodiversity and the restoration of many ecosystem functions when conditions become favorable [41, 42].

Following the rewetting incubation, the growth of several genera, including *Paenibacillus* and *Cohnella* from the phylum *Firmicutes*, were stimulated in more than half of soils (Fig. 3). Some genera within the *Firmicutes* (*Tumebacillus*, *Alicyclobacillus* and *Brevibacillus*) also showed the highest growth rate in many of the soils after rewetting. It is noted that all these genera belonged to order *Bacillales* within the *Firmicutes.* These endospore-forming bacteria thus represented the most active microbes or rapidly resuscitated organisms in these soils. Culture-based studies on a desert soil revealed growth of *Firmicutes* in cultivation medium, despite their low relative abundance in the soil [13]. Similarly, *Firmicutes* in many of our soils (S01, S03, S06, S09, S12 and S23) were not the most abundant group under desiccated conditions, but became dominant after the incubation (Fig. 2). These results suggested that the formation of endospores is not only beneficial for the survival of *Firmicutes* through water deprivation, but also an efficient strategy for recovery of populations when the environmental stress is relieved. Additionally, two genera within the *Gammaproteobacteria*, *Halomonas* and *Ralstonia*, also showed marked growth after the rewet (Fig. 3). Although members of *Halomonas* have not been reported to have strong resilience to extreme drought-rewet occurrences, they are halotolerant and capable of producing ectoine as osmolytes to maintain cell-osmotic balance [43]. The ectoine can protect the cells against salinity, desiccation and high temperature [44], which might facilitate cell survival in our soils following extreme moisture changes. Similarly, disaccharide trehalose contribute to osmotic stress tolerance of *Ralstonia* species [45], which might have implications for resilience to water stress.

It is noteworthy that *Actinobacteria* represent the second-most abundant group in desiccated soils (Fig. 2), suggesting high resistance to long-term desiccation as previously documented [13–15]. Some members of *Actinobacteria* contain genes for trace-gas scavenging, including H_2_ and CO, which can support a dormant lifestyle in arid deserts [15]. However, the growth rate of *Actinobacteria* appears much lower than *Firmicutes* and *Proteobacteria*, leading to sharp declines in average relative abundance in many of the soils after rewetting (Fig. 2). Similar trends were also found in a short period of a dry-rewet experiment, where the relative abundance of *Actinobacteria* increased after desiccation but then decreased following a rewet [30, 46]. It is also possible that the activity of *Actinobacteria* might be stimulated at the early stage of the 28-day incubation and outcompeted later by the stronger growth of *Firmicutes* and *Proteobacteria.* A previous study defined *Actinobacteria* as rapid responders with fast reactivation within one hour following rewetting, whereas *Firmicutes* and many *Proteobacteria* classes (*Alpha-*, *Beta-* and *Gamma-proteobacteria*) were consistently delayed responders [32]. This can be assessed by monitoring temporal changes in relative abundance of different phyla or classes in soils following rewetting in future studies.

Although all soil samples analyzed in the present study have been processed by the same laboratory treatment (long-term air-dry and storage in the same room conditions since 1939), it appeared that the microbial community compositions between different background soils can still be clearly distinguished, suggesting that the key bacteria identified in these soils should be indigenous to the soil field habitats rather than originating from the laboratory environment. This gained further support from analysis of some niche specialized taxa. For instance, members from class *Clostridia* (e.g. genus *Clostridium*), consisting of well-documented obligate anaerobes, were frequently detected (16.3-62.2%, Fig. S3a) in the desiccated rice paddy soils (S03 and S13-16), which should have endured constant anaerobic conditions by irrigation. In addition, although archaea were not detected or were only detected at low relative abundance in most of the soils, the class *Halobacteria*, known for salt tolerance, was exclusively detected in the two coastal saline soils (S22 and S23) in our study, accounting for >99% of the archaea (Fig. S4). These results suggest that genetic fingerprints of some microbes in desiccated soils is preserved (either as intact cell component or dead cell residues) and can be a used to indicate important field history, such as land use and soil formation. These results collectively validate the value of desiccated soil samples for systematic study on differences in soil microbial community composition, as previously proposed [18].

## 5 Conclusion

The present study revealed a diversified guild of bacteria resistant to seven decades of desiccation in 24 historical soils and nearly 25% of the taxa (at genus level) still show capability of growth. A dissimilar response of different phylotypes to the rewetting was observed, confirming to previous study suggesting the resuscitation strategies of soil microbes may be a phylogenetically conserved ecological trait [32]. Considering long-term starvation of microbes by desiccation and the fact that only water was added for incubation in our study, we expect growth of more diverse microbes with the additional supply of nutrients, such as carbon and nitrogen. Moreover, the activity of microbes was assessed by only DNA-based molecular analyses, which cannot capture all the active microbes in the soils, especially for those that did not quickly propagate within our incubation time. In addition, in future studies microbial viability would be more accurately depicted using more sensitive molecular tools, e.g. rRNA molecules rather than genes as a better indicator for cell activity [47]. Other methods would also be of great potential in distinguishing growing, dormant and dead cells, e. g. isotope tracing [31, 48–50] and single-cell based imaging [51, 52]. The removal of extracellular nucleic acids should also be considered in the future study to validate our conclusions [37]. Finally, future studies targeting specific microbial groups could enhance our understanding of some important functional microbes and their response to extreme stress [53]. By adopting more sensitive techniques, the desiccated historical soils could serve as precious “fossils” for soil microbiologists to discover tenacious species and retrieve undocumented site-specific historical events.

## Acknowledgements

We gratefully acknowledge the soil science predecessors of the Chinese Academy of Sciences for the original soil collection, the Institute of Soil Science for devotion to historical sample archives and all the former faculties in charge of the soil archiving. This work was supported by the National Science Foundation of China [41701302 and 41530857].

