## Supplementary for "Resuscitation of soil microbiota after > 70-years of desiccation"

[1]State Key Laboratory of Soil and Sustainable Agriculture. Institute of Soil Science, Chinese Academy of Sciences. Nanjing, 210008, Jiangsu Province, China

[2]University of Chinese Academy of Sciences, Beijing 100049, China

[3]School of Environmental Sciences, Norwich Research Park, University of East Anglia, Norwich, UK

*** Correspondence:**

Dongfeng Chen:

Marcela Hernández:

Running title: *Soil microbes after long-term desiccation*

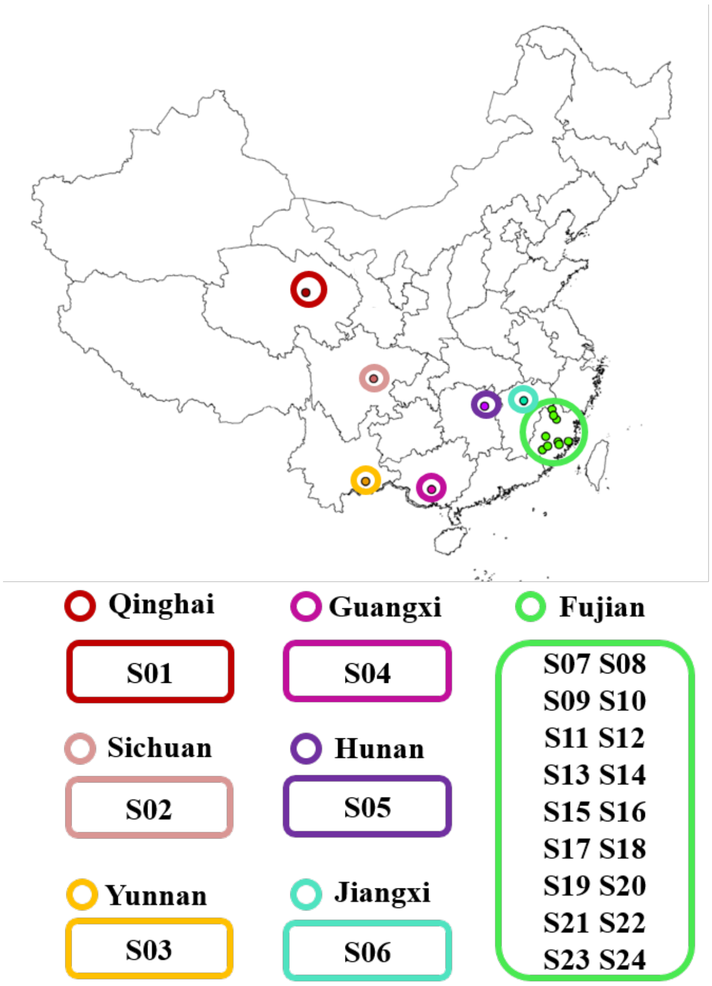


**Supplementary Figure 1.** The geographical distribution of soils collected between 1934-1939. The soils were collected from different districts across mainland of China.

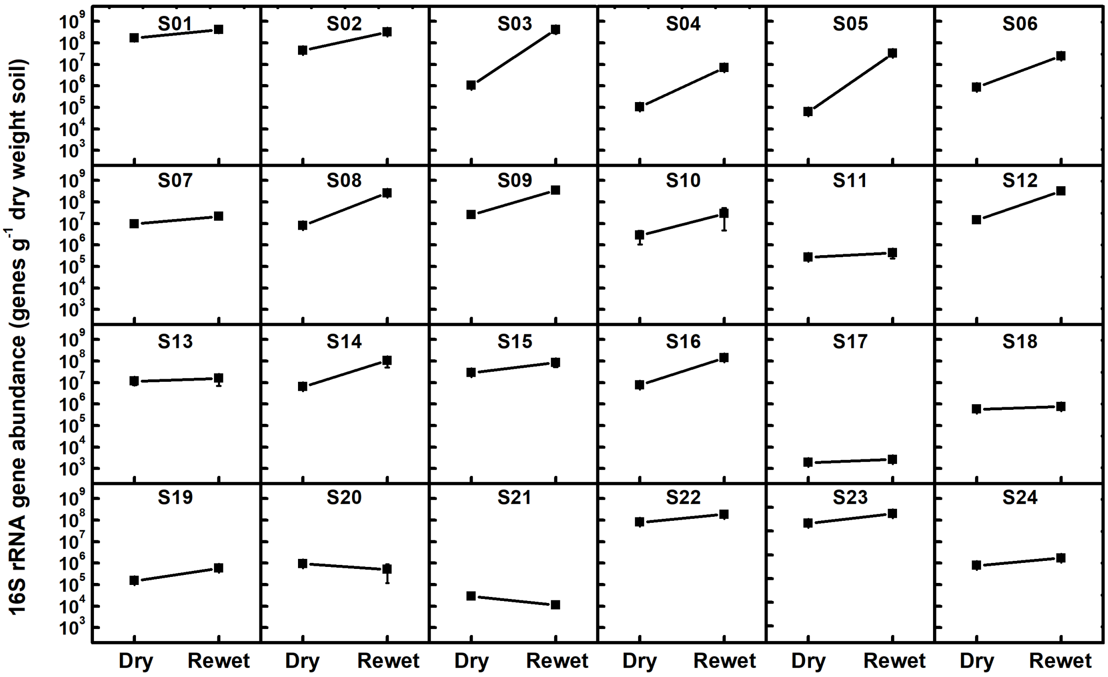


**Supplementary Figure 2.** Changes in bacterial abundance in 24 soils. The bacterial abundance was estimated at desiccated condition (Dry) and after rewetting incubation (Rewet) based on real-time quantitative PCR of 16S rRNA genes and high-throughput sequencing, and shown as mean value of 2-3 biological replicates with standard errors.

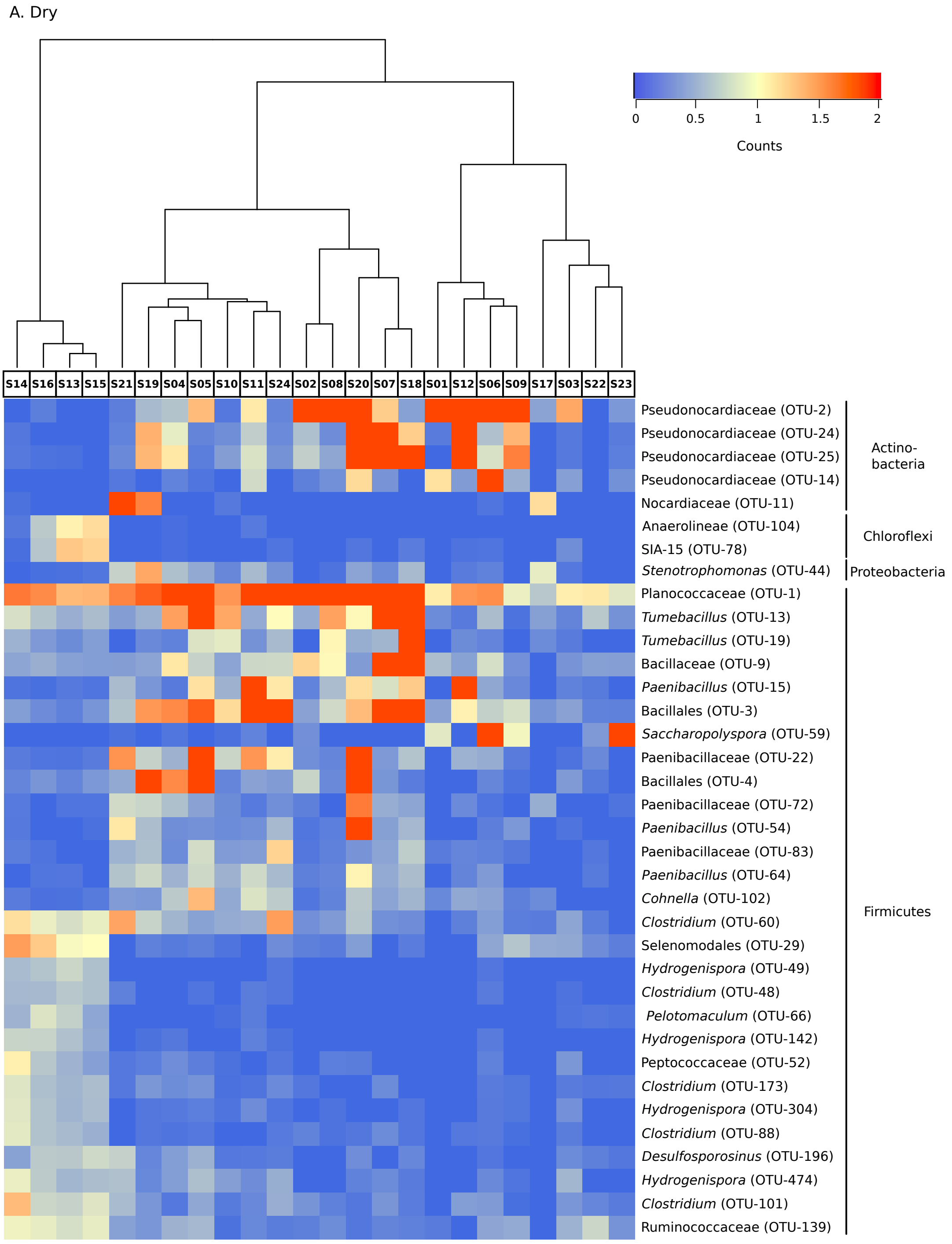


**Supplementary Figure 3.** Heatmap of the most relevant OTUs derived from total Bacteria. A. Dry soils; B. rewetted soils. The colored scale gives the percentage abundance of OTUs.

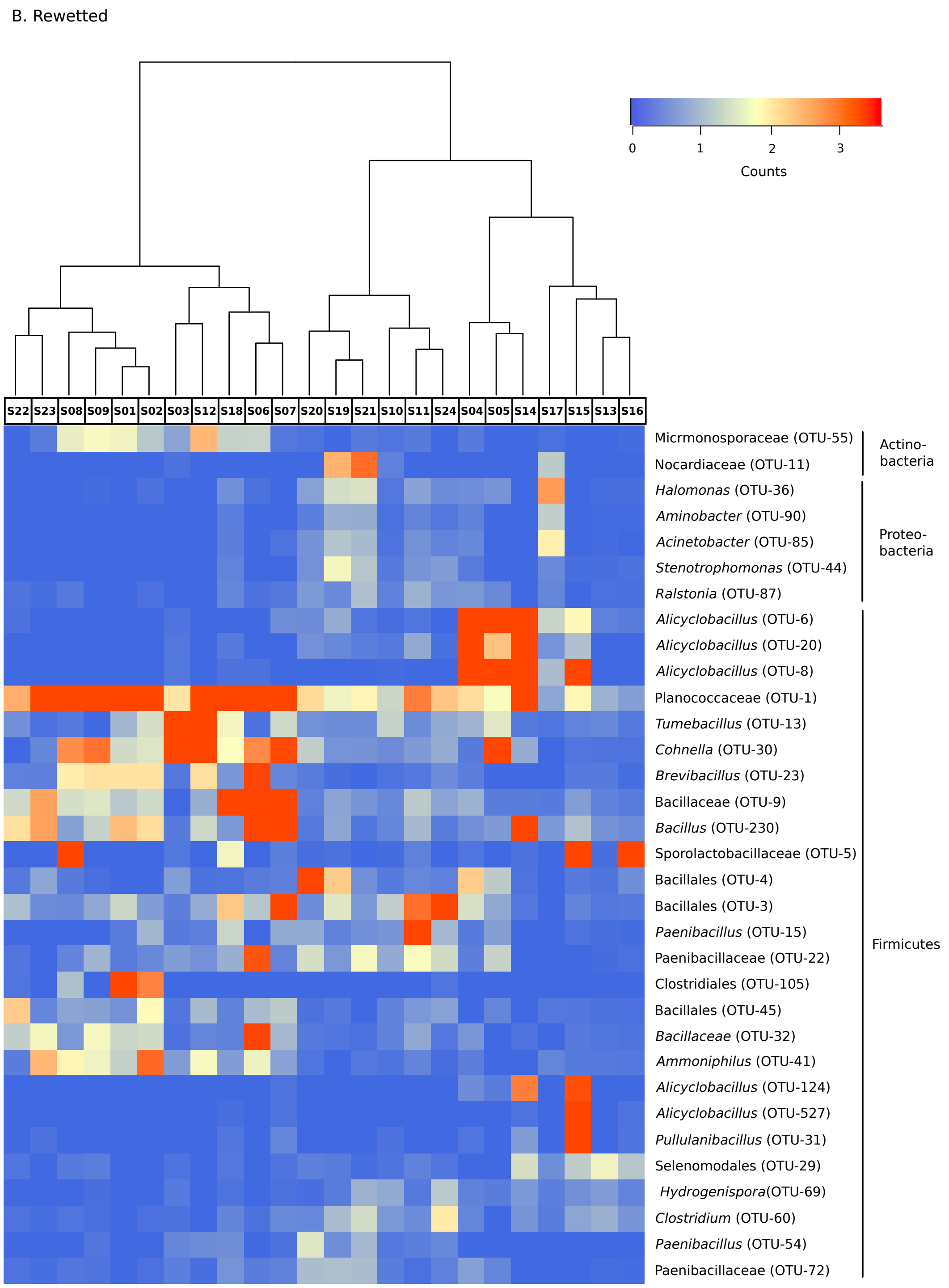


**Supplementary Figure 3. continues**

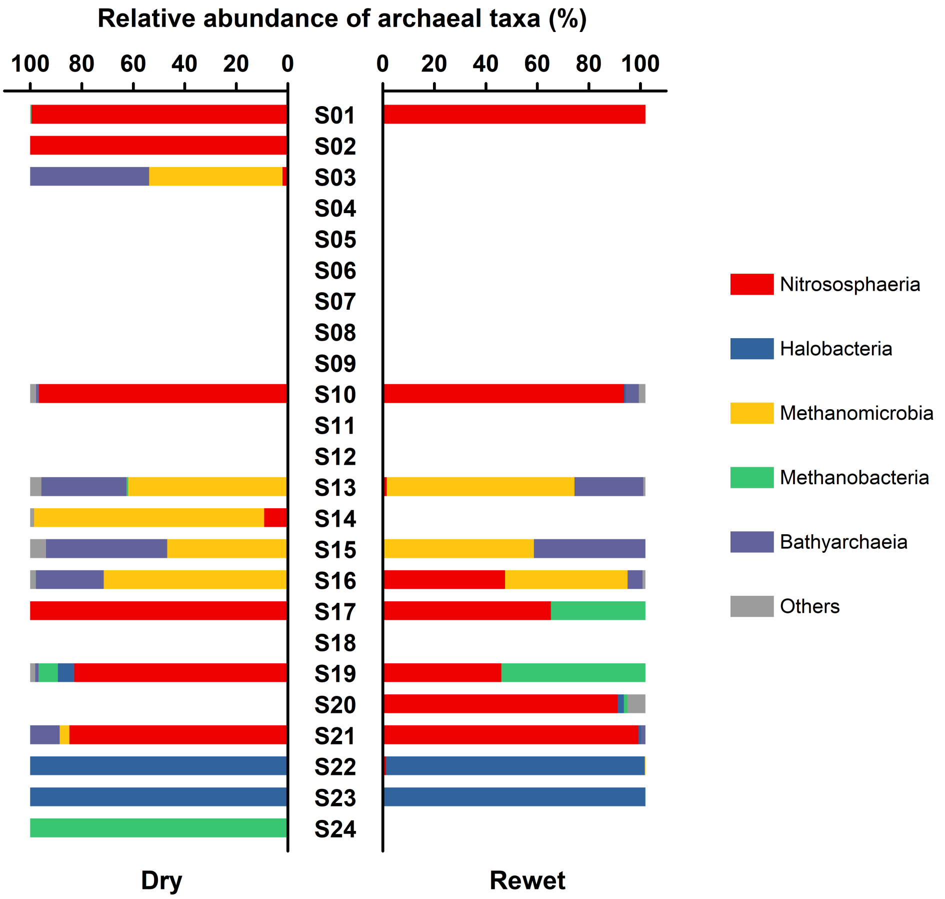


**Supplementary Figure 4.** Composition of the archaea in soils at desiccated condition (Dry) and after rewet incubation (Rewet). The archaeal composition was plotted at class level and the most abundant classes were shown. Taxa not seen more than 3 times in at least 20 % of the samples were removed using phyloseq package on R. The relative abundances of minor classes were summed up and shown as “others” which do not exceed 6.8% of total archaeal reads in any given soil.
